# Large eQTL meta-analysis reveals differing patterns between cerebral cortical and cerebellar brain regions

**DOI:** 10.1101/638544

**Authors:** Solveig K. Sieberts, Thanneer Perumal, Minerva M. Carrasquillo, Mariet Allen, Joseph S. Reddy, Gabriel E. Hoffman, Kristen K. Dang, John Calley, Philip J. Ebert, James Eddy, Xue Wang, Anna K. Greenwood, Sara Mostafavi, the AMP-AD Consortium, the CommonMind Consortium (CMC), Larsson Omberg, Mette Peters, Benjamin A. Logsdon, Philip L. De Jager, Nilüfer Ertekin-Taner, Lara M. Mangravite, AMP-AD Consortium, CommonMind Consortium

## Abstract

The availability of high-quality RNA-sequencing and genotyping data of post-mortem brain collections from consortia such as CommonMind Consortium (CMC) and the Accelerating Medicines Partnership for Alzheimer’s Disease (AMP-AD) Consortium enable the generation of a large-scale brain *cis*-eQTL meta-analysis. Here we generate cerebral cortical eQTL from 1433 samples available from four cohorts (identifying >4.1 million significant eQTL for >18,000 genes), as well as cerebellar eQTL from 261 samples (identifying 874,836 significant eQTL for >10,000 genes), and provide the results as a community resource. We find substantially improved power in the meta-analysis over individual cohort analyses, particularly in comparison to the Genotype-Tissue Expression (GTEx) Project eQTL. In addition, we observed differences in eQTL patterns between cerebral and cerebellar brain regions. We provide these brain eQTL as a common resource for use across the community in research programs. As a proof of principle for their utility, we apply a colocalization analysis to identify genes underlying the GWAS association peaks for schizophrenia and identify a potentially novel gene colocalization with lncRNA RP11-677M14.2 (posterior probability of colocalization 0.975).

## Introduction

Defining the landscape of genetic regulation of gene expression in a tissue-specific manner is useful for understanding both normal gene regulation and how variation in gene expression can alter disease risk. In the latter case, a variety of approaches now leverage the association between genetic variants and gene expression changes, including colocalization analysis^1–7^, transcriptome-wide association studies (TWAS)^8,9^, and gene regulatory network inference^10–16^.

There has been a relative lack of expression quantitative trait loci (eQTL) studies from the brain. Because of the more accessible nature of tissues such as blood or lymphoblastoid cell lines (LCLs), much of the large-scale identification of expression quantitative trait loci (eQTL) has occurred in these tissues^17–20^. For most other tissues, obtaining samples for RNA sequencing (RNA-seq) requires invasive biopsy, and brain tissues are typically only available in post-mortem brain samples. One effort, the Genotype-Tissue Expression (GTEx) project^21,22^, has profiled a broad range of tissues (42 distinct) for eQTL discovery, however, samples sizes in brain have been small (typically 100-150). Recently, efforts to understand gene expression changes in neuropsychiatric^23–25^ and neurodegenerative diseases^26–33^ have generated brain RNA-seq from disease and normal tissue, as well as genomewide genotypes. These analyses have found little evidence for widespread disease-specific eQTL, as well as high cross-cohort overlap^24,34^, meaning that most eQTL detected are disease-condition independent. This makes it possible to perform meta-analysis despite differences in disease ascertainment of the samples, to generate a well-powered brain eQTL resource for use in downstream research.

Here we generate a public eQTL resource from cerebral cortical tissue using 1433 samples from 4 cohorts from the CommonMind Consortium (CMC)^23,24^ and Accelerating Medicines Partnership for Alzheimer’s Disease (AMP-AD) Consortium^29,30^, as well as eQTL for cerebellum using 261 samples from AMP-AD. We show that eQTL discovered in GTEx, which consists of control individuals (without disease) only, are replicated in this larger brain eQTL resource. We further show widespread differences in regulation between cerebral cortex and cerebellum. To demonstrate one example of the utility of these data, we apply a colocalization analysis, which seeks to identify expression traits whose eQTL association pattern appears to co-occur at the same loci as the clinical trait association, to identify putative genes underlying the GWAS association peaks for schizophrenia^35^.

## Results

We generated eQTL from the publicly available AMP-AD (ROSMAP^26,27,34^ (Data Citation 1), Mayo RNAseq^28,36,37^ (Data Citation 2)) and CMC (MSSM-Penn-Pitt^24^ (Data Citation 3), HBCC (Data Citation 4)) cohorts with available genotypes and RNA-seq data, using a common analysis pipeline (Supplementary Table 1) (https://www.synapse.org/#!Synapse:syn17015233). Analyses proceeded separately by cohort. Briefly, the RNA-seq data were normalized for gene length and GC content prior to adjustment for clinical confounders, processing batch information, and hidden confounders using Surrogate Variable Analysis (SVA)^38^. Genes having at least 1 count per million (CPM) in at least 50% of samples were retained for downstream analysis (Supplementary Table 2). Genotypes were imputed to the Haplotype Reference Consortium (HRC) reference panel^39^. eQTL were generated adjusting for diagnosis (AD, control, other for AMP-AD cohorts and schizophrenia, control, bipolar/other for CMC cohorts) and principal components of ancestry separately for ROSMAP, Mayo temporal cortex (TCX), Mayo cerebellum (CER), MSSM-Penn-Pitt, and HBCC. For HBCC, which had a small number of samples derived from infant and adolescents, we excluded samples with age-of-death less than 18, to limit heterogeneity due to differences between the mature and developing brain.

We then performed a meta-analysis using the eQTL from cortical brain regions from the individual cohorts (dorsolateral prefrontal cortex (DLPFC) from ROSMAP, MSSM-Penn-Pitt, and HBCC and TCX from Mayo). The meta-analysis identifies substantially more eQTL than the individual cohorts (Table 1, Fig. 1). There is a strong relationship between the sample size in the individual cohorts and meta-analysis and the number of significant eQTL and genes with eQTL (Fig. 1b, 1c). Notably, the meta-analysis identified significant eQTL (at FDR ≤ 0.05) in >1000 genes for which no eQTL were observed in any individual cohort. Notably, we find significant eQTL for 18,295 (18,433 when considering markers with minor allele frequency (MAF) down to 1%) of the 19,392 genes included in the analysis.

**Figure 1:**
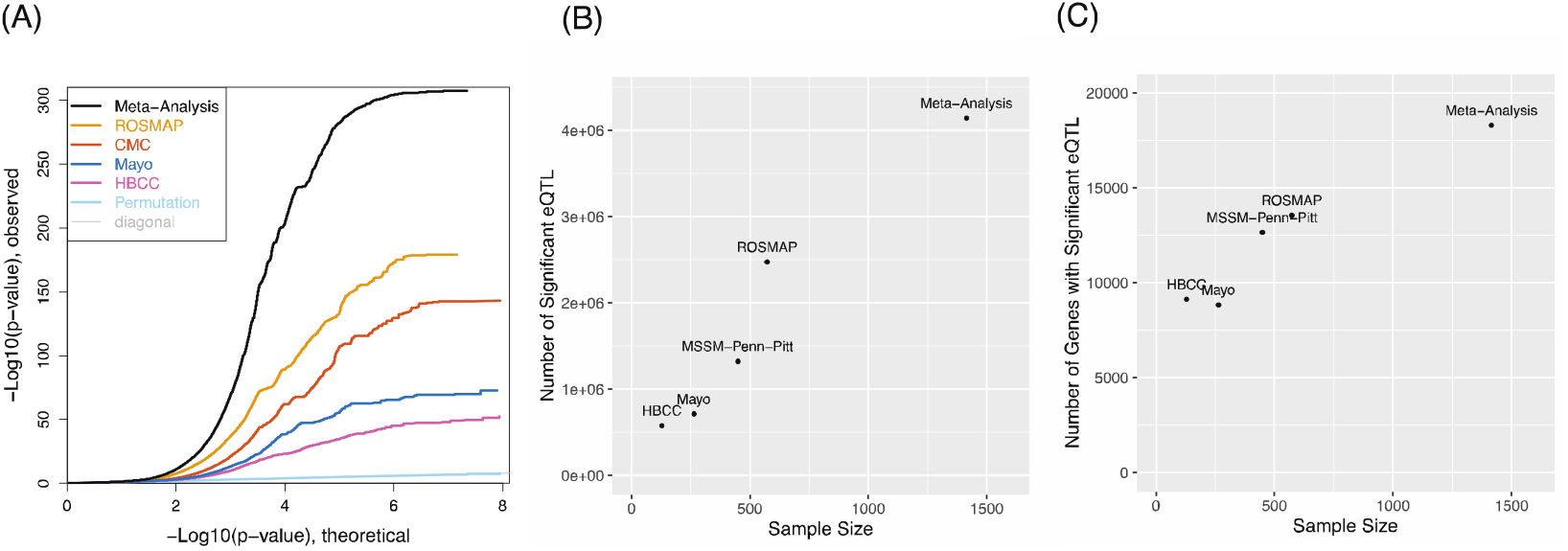
eQTL meta-analysis discovers more eQTL than individual cohorts. (A) Quantile-quantile plot of eQTL from individual cohorts as well as the meta-analysis of the true (black) and permuted (light blue) data. Number of significant eQTL (B) or genes with significant eQTL (C) as a function of cohort size.

We then compared our cortical eQTL to GTEx (v7)^21^, which is the most comprehensive brain eQTL database available in terms of number of available brain tissues (Table 1, Table 2). Due to the substantially larger power in these data, we find > 3.8 million eQTL not identified in GTEx cortical regions (Anterior Cingulate Cortex, Cortex or Frontal Cortex) and we find eQTL for >11,000 genes with no eQTL in these cortical regions in GTEx. We first evaluated the replication within our cortical and cerebellar eQTL of the region specific eQTL identified in GTEx. The cortical eQTL generated through the current analyses strongly replicate the eQTL available through GTEx, not only for cortical regions, but for all brain regions including cervical spinal cord (Table 2). Interestingly, the replication in these cortical eQTL of eQTL derived from the two GTEx cerebellar brain regions (cerebellum and cerebellar hemisphere) is consistently lower than for other brain regions represented in GTEx. However, replication of GTEx cerebellar eQTL is high when compared to the cerebellar eQTL generated in this analysis from the Mayo Clinic CER samples. We also performed the reverse comparison, by examining the replication of our eQTL in those region-specific eQTL identified in GTEx. Unsurprisingly, the replication levels were substantially lower, due to the lower power in the GTEx analyses.

**Table 1:** eQTL results from individual cohorts and meta-analysis.

| Cohort | Cis eQTL* | Unique Genes | eQTL (Genes) not Present in GTEx** | Meta-eQTL (Genes) not found in Cohort | Genes w/o eQTL in CMC, ROSMAP, HBCC or Mayo TCX |
| --- | --- | --- | --- | --- | --- |
| ROSMAP | 2,472,838 | 13,543 | 2,205,025 (7,829) | 2,103,711 (5,079) |  |
| Mayo TCX | 712,401 | 8,838 | 480,507 (4,439) | 3,485,381 (9,591) |  |
| CMC | 1,322,680 | 12,641 | 1,062,830 (7,206) | 2,948,886 (5,897) |  |
| HBCC | 577,512 | 9,112 | 379,732 (4,645) | 3,614,448 (9,337) |  |
| **Meta-Analysis\*\*\*** | **4,142,776** | **18,295** | **3,800,208 (11,395)** |  | **1,042** |

\* $MAF \geq 0.02$ and $FDR \leq 0.05$
\*\* GTEx (v7) Anterior Cingulate Cortex, Cortex or Frontal Cortex
\*\*\* Additional 36,352 (138) eQTL (Genes) with $0.01 \leq MAF < 0.02$

**Table 2:**
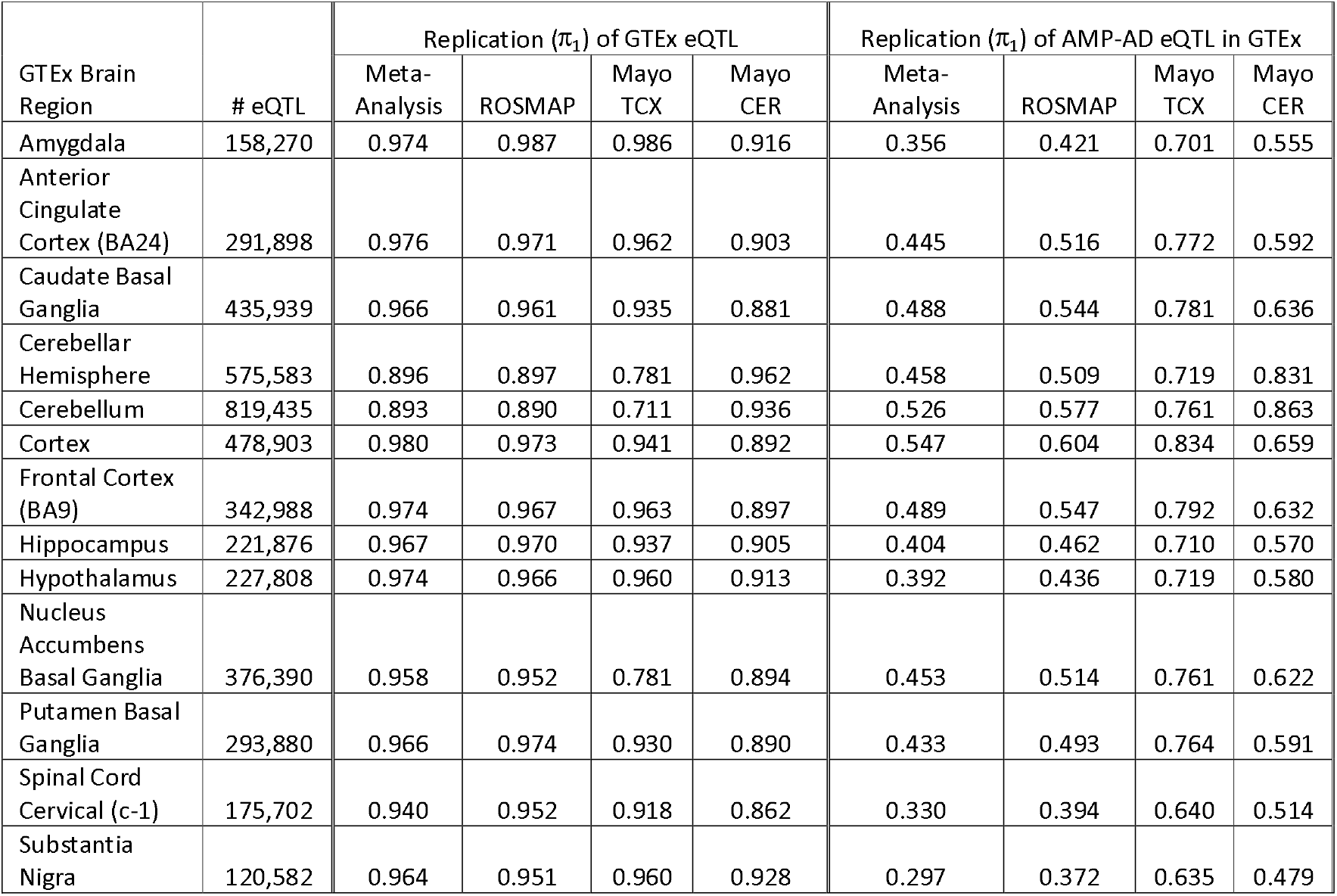
Replication rates between GTEx and publicly available eQTL from these analyses.

| GTEx Brain Region | # eQTL | Replication ($\pi_1$) of GTEx eQTL | | | | Replication ($\pi_1$) of AMP-AD eQTL in GTEx | | | |
| --- | --- | --- | --- | --- | --- | --- | --- | --- | --- |
|  |  | Meta-Analysis | ROSMAP | Mayo TCX | Mayo CER | Meta-Analysis | ROSMAP | Mayo TCX | Mayo CER |
| Amygdala | 158,270 | 0.974 | 0.987 | 0.986 | 0.916 | 0.356 | 0.421 | 0.701 | 0.555 |
| Anterior Cingulate Cortex (BA24) | 291,898 | 0.976 | 0.971 | 0.962 | 0.903 | 0.445 | 0.516 | 0.772 | 0.592 |
| Caudate Basal Ganglia | 435,939 | 0.966 | 0.961 | 0.935 | 0.881 | 0.488 | 0.544 | 0.781 | 0.636 |
| Cerebellar Hemisphere | 575,583 | 0.896 | 0.897 | 0.781 | 0.962 | 0.458 | 0.509 | 0.719 | 0.831 |
| Cerebellum | 819,435 | 0.893 | 0.890 | 0.711 | 0.936 | 0.526 | 0.577 | 0.761 | 0.863 |
| Cortex | 478,903 | 0.980 | 0.973 | 0.941 | 0.892 | 0.547 | 0.604 | 0.834 | 0.659 |
| Frontal Cortex (BA9) | 342,988 | 0.974 | 0.967 | 0.963 | 0.897 | 0.489 | 0.547 | 0.792 | 0.632 |
| Hippocampus | 221,876 | 0.967 | 0.970 | 0.937 | 0.905 | 0.404 | 0.462 | 0.710 | 0.570 |
| Hypothalamus | 227,808 | 0.974 | 0.966 | 0.960 | 0.913 | 0.392 | 0.436 | 0.719 | 0.580 |
| Nucleus Accumbens Basal Ganglia | 376,390 | 0.958 | 0.952 | 0.781 | 0.894 | 0.453 | 0.514 | 0.761 | 0.622 |
| Putamen Basal Ganglia | 293,880 | 0.966 | 0.974 | 0.930 | 0.890 | 0.433 | 0.493 | 0.764 | 0.591 |
| Spinal Cord Cervical (c-1) | 175,702 | 0.940 | 0.952 | 0.918 | 0.862 | 0.330 | 0.394 | 0.640 | 0.514 |
| Substantia Nigra | 120,582 | 0.964 | 0.951 | 0.960 | 0.928 | 0.297 | 0.372 | 0.635 | 0.479 |

Additionally, we compared our eQTL to a publically available fetal brain eQTL resource^40^ and found good replication of these eQTL as well (estimated replication rate π_1_ = 0.909 for the cortical meta-analysis, and π_1_ = 0.861 for cerebellum), though somewhat lower than the replication observed in the GTEx cohorts, which are comprised of adult-derived samples.

Finally, as a proof of concept, we performed a colocalization analysis between our eQTL meta-analysis and the Psychiatric Genomics Consortium (PGC) v2 schizophrenia GWAS summary statistics^35^. Seventeen genes showed posterior probability of colocalization using coloc^7^ (PP(H_4_)) >0.7 (Table 3), with 3 showing PP(H4) > 0.95 (FURIN, ZNF823, RP11-677M14.2). FURIN, having previously identified as a candidate through colocalization^24^ has recently been shown to reduce brain-derived neurotrophic factor (BDNF) maturation and secretion when inhibited by miR-338-3p^41^. ZNF823 has been identified in previous colocalization analyses^42,43^. RP11-677M14.2, a lncRNA located inside NRGN, while not previously identified through colocalization analysis, has been shown to be down-regulated in the amygdala of schizophrenia patients^44^. Noteably, NRGN does not appear to show eQTL colocalization (PP(H_4_) = 0.006), instead showing strong evidence for the eQTL and GWAS associations occurring independently (PP(H_3_) = 0.994).

**Table 3:**
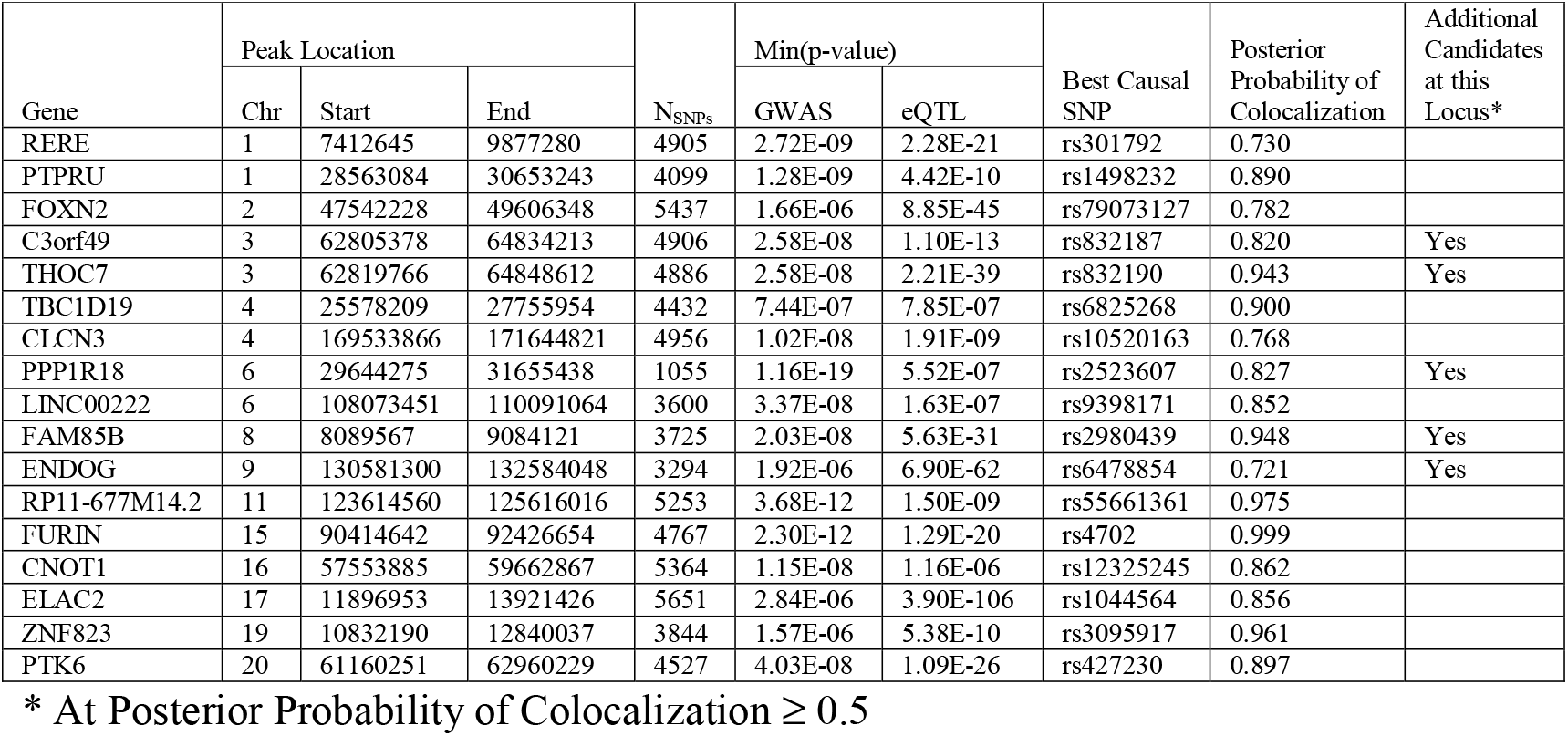
Top colocalized genes as inferred between meta-analysis eQTL and the PGC2 schizophrenia GWAS.

## Discussion

Using resources generated in the AMP-AD and CMC consortia, we have generated a well-powered brain eQTL resource for use by the scientific community. Unsurprisingly, we see a strong relationship between the number of significant eQTL, as well as genes with significant eQTL, and sample size using analyses from the individual cohorts and meta-analysis of those cohorts. This result has previously been shown for lower sample sizes^21^. We also show higher replication of GTEx eQTL in the meta-analysis relative to the individual cohorts.

Notably, we find significant eQTL for nearly every gene in our analysis, which include all but very lowly expressed genes (less than 1 cpm in more than 50% of samples). The wide discovery of eQTL is potentially beneficial for analyses utilizing these results, such as colocalization analysis or TWAS imputation, because more genes with significant eQTL means more genes can be evaluated with these approaches. Because we have discovered eQTL for most genes, further increasing sample size will not substantially increase the number of genes with significant eQTL, however it is likely that the number of significant eQTL associations within each gene would continue to increase, along with the accuracy of estimated effect sizes. This will result in a more accurate landscape of regulatory association, which will improve the ability to fine-map causal regions, and colocalize eQTL signal with clinical traits of interest. Thus, it will be valuable to continue to update this meta-analysis with additional data from these consortia and other resources as they become available, and continue to improve this resource as future data permits. Future work may also focus on using well-powered analysis to study the landscape of causal variation and covariation in gene regulation.

We found distinct eQTL patterns across cerebral cortical and cerebellar brain regions in our resource. Specifically, comparison of eQTL from our resource with those from GTEx shows high replication for the majority of brain regions. However, cerebellar regions show consistently lower replication with the cerebral cortical eQTL generated here. In contrast, the cerebellar eQTL generated from the Mayo Clinic study replicate GTEx cerebellar eQTL at a substantially higher rate, suggesting a different pattern of regulatory variation affecting expression in cerebellum versus other brain regions. Indeed, epigenomic analyses show substantial differences between cerebellar and cerebral cortical regions^45–48^, particularly in methylation patterns, which could drive different eQTL association patterns. This is further corroborated by the observation of substantial coexpression differences between cerebellar and other brain regions^49^. These effects could be due to differences in cell type composition, with cerebellar regions consisting of substantially more neurons than other brain regions^50^. This is supported by a gene enrichment analysis of genes showing cerebellum-specific eQTL patterns, which showed that most of the top gene sets were related to axon and neuron generation and differentiation (Supplementary Table 3). One recent report suggests that there are also widespread differences in histone modifications within cell types derived from cerebellar and cortical regions^51^, though this effect had not been noted in other studies. In particular, Ma et al^51^ observed that both neuronal and non-neuronal cell types show differing histone modifications across tissue of origin. Further work is necessary to confirm this finding and to develop models to deconvolve the cell-type specific regulatory effects in different brain regions^52–54^, however our analysis demonstrates that this meta-analysis is representative of eQTL across the majority of brain regions, with the exception of cerebellum. Future meta-analytic analyses may also cast a wider net in terms of brain regions included.

The replication of fetal eQTL, while significant, is somewhat lower than the replication of adult eQTL represented in GTEx. This may be due to multiple factors. The fetal eQTL analysis was generated from brain homogenate, rather than dissected brain regions, though the lower replication likely also reflects broad transcriptional differences between developing and mature brain^55^. These transcriptional differences may also explain why we find substantially more eQTL than a recently published, similarly sized eQTL analysis which uses samples from across developmental and adult timepoints^25^, and why this meta-analysis shows higher replication of GTEx eQTL.

Finally, the lack of widespread disease-specific eQTL observed in schizophrenia (CMC)^24^ and Alzheimer’s (ROSMAP)^34^, as well as a strong overlap among eQTL derived from samples from individuals with these diseases, as well as normal individuals from these and other cohorts such as GTEx^24,34^, suggests that disease-specific eQTL, if they exist, are likely few in number and/or small in effect size. Thus, the heterogeneous samples derived from different disease-based cohorts can be meta-analyzed to create a general-purpose brain eQTL resource representing adult gene regulation, despite comprising samples with different disease backgrounds, along with normal controls. Therefore, these eQTL will be useful both within and outside these specific disease contexts. For example, since these eQTL are not disease specific they may be used to understand healthy gene expression regulation in the brain, as well as to infer colocalization of eQTL signatures with disease risk for any disease whose tissue etiology is from the brain, since these signatures are reflective of normal brain regulation. It should be stated that while many eQTL are not disease specific, i.e. they are identified under various central nervous system (CNS) disease diagnoses and in control brains, they may still contribute to common CNS diseases as previously demonstrated^24,31–33,42,43^. While we have demonstrated a proof-of-concept colocalization analysis with a previously published schizophrenia GWAS, these eQTL are a broadly useful resource for studying neuropsychiatric and neurodegenerative disorders, as well as for understanding the landscape of gene regulation in brain.

## Methods

### RNA-seq Re-alignment

For the CMC studies (MSSM-Penn-Pitt, HBCC), RNA-seq reads were aligned to GRCh37 with STAR v2.4.0g1 ^56^ from the original FASTQ files. Uniquely mapping reads overlapping genes were counted with featureCounts v1.5.2^57^ using annotations from ENSEMBL v75.

For the AMP-AD studies (ROSMAP, Mayo RNAseq), Picard v2.2.4 (https://broadinstitute.github.io/picard/) was used to generate FASTQ files from the available BAM files, using the Picard SamToFastq function. Picard SortSam was first applied to ensure that R1 and R2 reads were correctly ordered in the intermediate SAM file before converting to FASTQ. The converted FASTQs were aligned to the GENCODE24 (GRCh38) reference genome using STAR v2.5.1b, with twopassMode set as Basic. Gene counts were computed for each sample by STAR by setting quantMode as GeneCounts.

### RNA-seq Normalization

To account for differences between samples, studies, experimental batch effects and unwanted RNA-seq-specific technical variations, we performed library normalization and covariate adjustments for each study separately using fixed/mixed effects modeling. The workflow consisted of following steps:

1. ***Gene filtering:*** Out of ~56K aligned and quantified genes only genes showing at least modest expression were used in this analysis. Genes that were expressed more than 1 CPM (read Counts Per Million total reads) in at least 50% of samples in each tissue and diagnosis category was retained for analysis. Additionally, genes with available gene length and percentage GC content from BioMart December 2016 archive were subselected from the above list. This resulted in approximately 14K to 16K genes in each study.
2. ***Calculation of normalized expression values:*** Sequencing reads were then normalized in two steps. First, conditional quantile normalization (CQN)^58^ was applied to account for variations in gene length and GC content. In the second step, the confidence of sampling abundance was estimated using a weighted linear model using voom-limma package in bioconductor^59,60^. The normalized observed read counts, along with the corresponding weights, were used in the following steps.
3. ***Outlier detection:*** Based on normalized log2(CPM) of expression values, outlier samples were detected using principal component analysis (PCA)^61,62^ and hierarchical clustering. Samples identified as outliers using both the above methods were removed from further analysis.
4. ***Covariate imputation:*** Before identifying associated covariates, important missing covariates were imputed. Principally, post-mortem interval (PMI), or the latency between death and tissue collection, which is frequently an important covariate for the analysis of gene expression from post-mortem brain tissue, was imputed for a portion of samples in Mayo RNAseq data for which true values were unavailable. Genomic predictors of PMI were estimated using ROSMAP and MSSM (an additional RNA-seq study available through AMP-AD) samples and were used to impute missing values as necessary.
5. ***Covariate identification:*** Normalized log2(CPM) counts were then explored to determine which known covariates (both biological and technical) should be adjusted. Except for the HBCC study, we used a stepwise (weighted) fixed/mixed effect regression modeling approach to select the relevant covariates having a significant association with gene expression. Here, covariates were sequentially added to the model if they were significantly associated with any of the top principal components, explaining more than 1% of variance of expression residuals. For HBCC, we used a model selection based on Bayesian information criteria (BIC) to identify the covariates that improve the model in a greater number of genes than making it worse.
6. ***SVA adjustments:*** After identifying the relevant known confounders, hidden-confounders were identified using the Surrogate Variable Analysis (SVA)^38^. We used a similar approach as previously defined^24^ to find the number of surrogate variables (SVs), which is more conservative than the default method provided by the SVA package in R^63^. The basic idea of this approach is that for an eigenvector decomposition of permuted residuals each eigenvalue should explain an equal amount of the variation. By the nature of eigenvalues, however, there will always be at least one that exceeds the expected value. Thus, from a series of 100 permutations of residuals (white noise) we identified the number of covariates as shown in Supplementary Table 1. We applied the “irw” (iterative re-weighting) version of SVA to the normalized gene expression matrix, along with the covariate model described above to obtain residual gene expression for eQTL analysis.
7. ***Covariate adjustments:*** We performed a variant of fixed/mixed effect linear regression, choosing mixed-effect models when multiple tissues or samples, were available per individual, as shown here: gene expression ~ Diagnosis + Sex + covariates + (1|Donor), where each gene in linearly regressed independently on Diagnosis, identified covariates and donor (individual) information as random effect. Observation weights (if any) were calculated using the voom-limma^59,60^ pipeline, which has a net effect of up-weighting observations with inferred higher precision in the linear model fitting process to adjust for the mean-variance relationship in RNA-seq data. The Diagnosis component was then added back to the residuals to generate covariate-adjusted expression for eQTL analysis.

All these workflows were applied separately for each study. For the AMP-AD studies, gene locations were lifted over to GRCh37 for comparison with the genotype imputation panel (described below). For HBCC, samples with age < 18 were excluded prior to analysis. Supplementary Table 1 shows the covariates and surrogate variables identified in each study.

### AD Diagnosis Harmonization

Prior to RNA-seq normalization, we harmonized the LOAD definition across AMP-AD studies. AD controls were defined as patients with a low burden of plaques and tangles, as well as lack of evidence of cognitive impairment. For the ROSMAP study, we defined AD cases to be individuals with a Braak^64^ greater than or equal to 4, CERAD score^65^ less than or equal to 2, and a cognitive diagnosis of probable AD with no other causes (cogdx=4), and controls to be individuals with Braak less than or equal to 3, CERAD score greater than or equal to 3, and cognitive diagnosis of ‘no cognitive impairment’ (cogdx = 1). For the Mayo Clinic study, we defined disease status based on neuropathology, where individuals with Braak score greater than or equal to 4 were defined to be AD cases, and individuals with Braak less than or equal to 3 were defined to be controls. Individuals not meeting “AD case” or “control” criteria were retained for analysis, and were categorized as “other” for the purposes of RNA-seq adjustment.

### Genotype QC and Imputation

Genotype QC was performed using PLINK v1.9^66^. Markers with zero alternate alleles, genotyping call rate ≤ 0.98, Hardy-Weinberg *p*-value < 5e-5 were removed, as well as individuals with genotyping call rate < 0.90. Samples were then imputed to HRC (Version r1.1 2016)^39^, as follows: if necessary marker positions were lifted-over to GRCh37 and aligned to the HRC loci using HRC-1000G-check-bim-v4.2 (http://www.well.ox.ac.uk/~wrayner/tools/), which checks the strand, alleles, position, reference/alternate allele assignments and frequencies of the markers, removing A/T & G/C single nucleotide polymorphisms (SNPs) with minor allele frequency (MAF) > 0.4, SNPs with differing alleles, SNPs with > 0.2 allele frequency difference between the genotyped samples and the HRC samples, and SNPs not in reference panel. Imputation was performed via the Michigan Imputation Server^67^ using Eagle v2.3^68^ as the phasing algorithm. Imputation was done separately by cohort and by chip within cohort, and markers with R^2^ ≥ 0.7 and minor allele frequency (MAF) ≥ 0.01 (within cohort) were retained for analysis.

### Genetic Ancestry Inference

GEMTOOLs^69^ was used to infer ancestry and compute ancestry components separately by cohort. The number of significant ancestry components were also estimated by the GEMTOOLs algorithm. For MSSM-Penn-Pitt and HBCC, which are multi-ethnic cohorts, only Caucasian samples were retained for eQTL analysis.

### eQTL Analysis

eQTL were generated separately in each cohort using MatrixEQTL^70^ adjusting for harmonized Diagnosis and inferred Ancestry components using “cis” gene-marker comparisons: Expression ~ Genotype + Diagnosis + PC_1_ + … + PC_n,_, where PC_k_ is the k^th^ ancestry component, using Expression variables which were previously covariate adjusted as described above. Here we define “cis” as ± 1 MB around the gene, and GRCh37 gene locations were used for consistency with the marker imputation panel. Meta-analysis was performed via fixed-effect model^71^ using an adaptation of the metareg function in the gap package in R. Given that multiple tissues were present, we also evaluated a random-effect model, but found it to be highly conservative in this case. In order to assess potential inflation of Type 1 error, we performed 5 permutations of this analysis, starting by permuting gene expression, relative to genotype and ancestry components, within diagnosis for each cohort, and performing meta-analysis of the permuted eQTL. We found that Type 1 error was well controlled (Fig. 1a).

### Comparison with GTEx and Fetal eQTL

Full summary statistics for the GTEx v7^21^ eQTL for all available brain regions were obtained from the GTEx Portal (https://gtexportal.org/), and fetal eQTL were obtained from Figshare^72^. Markers and genes present in the external eQTL as well as our analysis were retained for comparison. The replication rate was estimated as the π_1_ statistic using the qvalue package^73^ in R as follows: the meta-analysis p-values for significant (at FDR 0.05) GTEx were used to estimate the replication rate of GTEx eQTL in the meta-analysis. Analogous methods were used to estimate all other replication rates.

### Pathway Analysis of Cerebellar eQTL Genes

In order to identify whether genes showing cerebellar-specific eQTL patterns showed any biological coherence, we performed a pathway analysis as follows. For genes with at least 5 significant cerebellar eQTL, we computed the correlation of effect-size between cerebellum eQTL and cortical eQTL for the loci that were significant in cerebellum. We then selected genes for which the effect-sizes were uncorrelated (*p*-value > 0.05) between the two tissues as showing cerebellar-specific eQTL patterns, and performed a pathway analysis with GO biological processes, cellular components and molecular function using a Fisher’s exact test. Note that due to the (power-mediated) greater detection of eQTL in cortex, we did not perform the reverse comparison.

### Coloc Analysis

We applied Approximate Bayes Factor colocalization (coloc.abf)^7^ from the coloc R package to the summary statistics from the PGC2 Schizophrenia GWAS^35^ downloaded from the PGC website (http://pgc.unc.edu), and the summary statistics from the eQTL meta-analysis. Each gene present in the meta-analysis was compared to the GWAS in turn, and suggestive and significant GWAS peaks with *p*-value < 5e-6 were considered for analysis.

### Code availability

An R package with all code for the gene expression normalization is available at https://github.com/Sage-Bionetworks/ampad-diffexp. All other analyses were generated using packages publically available from their respective authors.

## Data Records

eQTL results for the ROSMAP, Mayo TCX, Mayo CER and cortical meta-analysis are found in Data Citation 5, Data Citation 6, Data Citation 7 and Data Citation 8, respectively, in the AMP-AD Knowledge Portal. These results include SNP (location, rsid, alleles, and allele frequency) and gene (location, gene symbol, strand and biotype) information, as well as estimated effect size (beta), statistic (z), p-value, FDR, and expression-increasing allele.

## Acknowledgements

For the ROSMAP and Mayo RNAseq studies, the results published here are in whole or in part based on data obtained from the AMP-AD Knowledge Portal (doi:10.7303/syn2580853). ROSMAP study data were provided by the Rush Alzheimer’s Disease Center, Rush University Medical Center, Chicago. Data collection was supported through funding by NIA grants P30AG10161, R01AG15819, R01AG17917, R01AG30146, R01AG36836, U01AG32984, U01AG46152, the Illinois Department of Public Health, and the Translational Genomics Research Institute. Mayo RNA-seq study data were provided by the following sources: The Mayo Clinic Alzheimers Disease Genetic Studies, led by Dr. Nilufer Ertekin-Taner and Dr. Steven G. Younkin, Mayo Clinic, Jacksonville, FL using samples from the Mayo Clinic Study of Aging, the Mayo Clinic Alzheimer’s Disease Research Center, and the Mayo Clinic Brain Bank. Data collection was supported through funding by NIA grants P50 AG016574, R01 AG032990, U01 AG046139, R01 AG018023, U01 AG006576, U01 AG006786, R01 AG025711, R01 AG017216, R01 AG003949, NINDS grant R01 NS080820, CurePSP Foundation, and support from Mayo Foundation. Study data includes samples collected through the Sun Health Research Institute Brain and Body Donation Program of Sun City, Arizona. The Brain and Body Donation Program is supported by the National Institute of Neurological Disorders and Stroke (U24 NS072026 National Brain and Tissue Resource for Parkinsons Disease and Related Disorders), the National Institute on Aging (P30 AG19610 Arizona Alzheimers Disease Core Center), the Arizona Department of Health Services (contract 211002, Arizona Alzheimers Research Center), the Arizona Biomedical Research Commission (contracts 4001, 0011, 05-901 and 1001 to the Arizona Parkinson’s Disease Consortium) and the Michael J. Fox Foundation for Parkinsons Research. This study was in part supported by NIH RF1 AG051504 and R01 AG061796 (NET).

For CommonMind, data were generated as part of the CommonMind Consortium supported by funding from Takeda Pharmaceuticals Company Limited, F. Hoffmann-La Roche Ltd and NIH grants R01MH085542, R01MH093725, P50MH066392, P50MH080405, R01MH097276, RO1-MH-075916, P50M096891, P50MH084053S1, R37MH057881, AG02219, AG05138, MH06692, R01MH110921, R01MH109677, R01MH109897, U01MH103392, and contract HHSN271201300031C through IRP NIMH. Brain tissue for the study was obtained from the following brain bank collections: the Mount Sinai NIH Brain and Tissue Repository, the University of Pennsylvania Alzheimer’s Disease Core Center, the University of Pittsburgh NeuroBioBank and Brain and Tissue Repositories, and the NIMH Human Brain Collection Core. CMC Leadership: Panos Roussos, Joseph Buxbaum, Andrew Chess, Schahram Akbarian, Vahram Haroutunian (Icahn School of Medicine at Mount Sinai), Bernie Devlin, David Lewis (University of Pittsburgh), Raquel Gur, Chang-Gyu Hahn (University of Pennsylvania), Enrico Domenici (University of Trento), Mette A. Peters, Solveig Sieberts (Sage Bionetworks), Thomas Lehner, Geetha Senthil, Stefano Marenco, Barbara K. Lipska (NIMH).

SKS, TP, KKD, JE, AKG, LO, BAL, and LMM were additionally supported by NIA grants U24 AG61340, U01 AG46170, U01 AG 46161, R01 AG46171, R01 AG 46174.

All data used in this manuscript have been previously released through their respective consortia, and have been reviewed by IRBs at their institution of origin. Informed consent has been obtained from all individuals.

## Author contributions

SKS, SM, MP, PLDJ, NET, LMM, the AMP-AD Consortium, and the CMC Consortium contributed to the design and generation of the study data. SKS, TP, MC, MA, JSR, GH, KDD, JC, PJE, JE contributed the data analysis. SKS, TP, GH, AKG, LO, MP, BAL, LMM contributed to the manuscript preparation and data sharing.

## Data Citations

1. *Synapse* (ROSMAP) 10.7303/syn3219045
2. *Synapse* (Mayo) 10.7303/syn5550404
3. *Synapse* (MSSM-Penn-Pitt) 10.7303/syn2759792
4. *Synapse* (HBCC) 10.7303/syn2759792
5. *Synapse* (ROSMAP eQTL) 10.7303/syn16984409.1
6. *Synapse* (Mayo TCX eQTL) 10.7303/syn16984410.1
7. *Synapse* (Mayo CER eQTL) 10.7303/syn16984411.1
8. *Synapse* (meta-analysis eQTL) 10.7303/syn16984815.1

## References

1. Zhu, Z. et al. Integration of summary data from GWAS and eQTL studies predicts complex trait gene targets. Nat. Genet. 48, 481–487 (2016).

2. Nica, A. C. et al. Candidate Causal Regulatory Effects by Integration of Expression QTLs with Complex Trait Genetic Associations. PLoS Genet. 6, e1000895 (2010).

3. Ongen, H. et al. Estimating the causal tissues for complex traits and diseases. Nat. Genet. 49, 1676–1683 (2017).

4. Chun, S. et al. Limited statistical evidence for shared genetic effects of eQTLs and autoimmune-disease-associated loci in three major immune-cell types. Nat. Genet. 49, 600–605 (2017).

5. Hormozdiari, F., Kostem, E., Kang, E. Y., Pasaniuc, B. & Eskin, E. Identifying Causal Variants at Loci with Multiple Signals of Association. Genetics 198, 497–508 (2014).

6. He, X. et al. Sherlock: Detecting Gene-Disease Associations by Matching Patterns of Expression QTL and GWAS. Am. J. Hum. Genet. 92, 667–680 (2013).

7. Giambartolomei, C. et al. Bayesian Test for Colocalisation between Pairs of Genetic Association Studies Using Summary Statistics. PLoS Genet. 10, e1004383 (2014).

8. Gamazon, E. R. et al. A gene-based association method for mapping traits using reference transcriptome data. Nat. Genet. 47, 1091–1098 (2015).

9. Barbeira, A. N. et al. Exploring the phenotypic consequences of tissue specific gene expression variation inferred from GWAS summary statistics. bioRxiv 45260 (2017). doi:10.1101/045260

10. Zhu, J. et al. An integrative genomics approach to the reconstruction of gene networks in segregating populations. Cytogenet. Genome Res. 105, 363–74 (2004).

11. Schadt, E. et al. Mapping the Genetic Architecture of Gene Expression in Human Liver. PLoS Biol. 6, e107 (2008).

12. Zhu, J. et al. Integrating large-scale functional genomic data to dissect the complexity of yeast regulatory networks. Nat. Genet. 40, 854–61 (2008).

13. Greenawalt, D. M. et al. A survey of the genetics of stomach, liver, and adipose gene expression from a morbidly obese cohort. Genome Res. 21, (2011).

14. Zhang, B. et al. Integrated systems approach identifies genetic nodes and networks in late-onset Alzheimer’s disease. Cell 153, 707–20 (2013).

15. Franzén, O. et al. Cardiometabolic risk loci share downstream cis- and transgene regulation across tissues and diseases. Science 353, 827–30 (2016).

16. Peters, L. A. et al. A functional genomics predictive network model identifies regulators of inflammatory bowel disease. Nat. Genet. 49, 1437–1449 (2017).

17. Battle, A. et al. Characterizing the genetic basis of transcriptome diversity through RNA-sequencing of 922 individuals. Genome Res. 24, 14–24 (2014).

18. Westra, H.-J. et al. Systematic identification of trans eQTLs as putative drivers of known disease associations. Nat. Genet. 45, 1238–1243 (2013).

19. Võsa, U. et al. Unraveling the polygenic architecture of complex traits using blood eQTL meta-analysis. bioRxiv 447367 (2018). doi:10.1101/447367

20. Qi, T. et al. Identifying gene targets for brain-related traits using transcriptomic and methylomic data from blood. Nat. Commun. 9, 2282 (2018).

21. Aguet, F. et al. Genetic effects on gene expression across human tissues. Nature 550, 204–213 (2017).

22. Ardlie, K. G. et al. The Genotype-Tissue Expression (GTEx) pilot analysis: Multitissue gene regulation in humans. Science (80-.). 348, 648–660 (2015).

23. CommonMind Consortium Knowledge Portal. (2014). doi:10.7303/syn2759792

24. Fromer, M. et al. Gene expression elucidates functional impact of polygenic risk for schizophrenia. Nature Neuroscience 19, 1442–1453 (2016).

25. Wang, D. et al. Comprehensive functional genomic resource and integrative model for the human brain. Science (80-.). 362, eaat8464 (2018).

26. Chibnik, L. B. et al. Susceptibility to neurofibrillary tangles: role of the PTPRD locus and limited pleiotropy with other neuropathologies. Mol. Psychiatry 23, 1521 (2017).

27. Mostafavi, S. et al. A molecular network of the aging human brain provides insights into the pathology and cognitive decline of Alzheimer’s disease. Nat. Neurosci. 21, 811–819 (2018).

28. Allen, M. et al. Human whole genome genotype and transcriptome data for Alzheimer’s and other neurodegenerative diseases. Sci. Data 3, 160089 (2016).

29. Logsdon, B. et al. Meta-analysis of the human brain transcriptome identifies heterogeneity across human AD coexpression modules robust to sample collection and methodological approach. bioRxiv 510420 (2019). doi:10.1101/510420

30. AMP-AD Knowledge Portal. (2015). doi:10.7303/syn2580853

31. Allen, M. et al. Novel late-onset Alzheimer disease loci variants associate with brain gene expression. Neurology 79, 221–8 (2012).

32. Zou, F. et al. Brain expression genome-wide association study (eGWAS) identifies human disease-associated variants. PLoS Genet. 8, e1002707 (2012).

33. Allen, M. et al. Late-onset Alzheimer disease risk variants mark brain regulatory loci. Neurol. Genet. 1, e15 (2015).

34. Ng, B. et al. An xQTL map integrates the genetic architecture of the human brain’s transcriptome and epigenome. Nat. Neurosci. 20, 1418–1426 (2017).

35. Ripke, S. et al. Biological insights from 108 schizophrenia-associated genetic loci. Nature 511, 421–427 (2014).

36. Allen, M. et al. Conserved brain myelination networks are altered in Alzheimer’s and other neurodegenerative diseases. Alzheimers. Dement. 14, 352–366 (2018).

37. Allen, M. et al. Divergent brain gene expression patterns associate with distinct cell-specific tau neuropathology traits in progressive supranuclear palsy. Acta Neuropathol. 136, 709–727 (2018).

38. Leek, J. T. & Storey, J. D. Capturing Heterogeneity in Gene Expression Studies by Surrogate Variable Analysis. PLoS Genet. 3, e161 (2007).

39. McCarthy, S. et al. A reference panel of 64,976 haplotypes for genotype imputation. Nat. Genet. 48, 1279–83 (2016).

40. O’Brien, H. E. et al. Expression quantitative trait loci in the developing human brain and their enrichment in neuropsychiatric disorders. Genome Biol. 19, 194 (2018).

41. Hou, Y. et al. Schizophrenia-associated rs4702 G allele-specific downregulation of FURIN expression by miR-338-3p reduces BDNF production. Schizophr. Res. 199, 176–180 (2018).

42. Dobbyn, A. et al. Landscape of Conditional eQTL in Dorsolateral Prefrontal Cortex and Co-localization with Schizophrenia GWAS. Am. J. Hum. Genet. 102, 1169–1184 (2018).

43. Pardiñas, A. F. et al. Common schizophrenia alleles are enriched in mutation-intolerant genes and in regions under strong background selection. Nat. Genet. 50, 381–389 (2018).

44. Liu, Y. et al. Non-coding RNA dysregulation in the amygdala region of schizophrenia patients contributes to the pathogenesis of the disease. Transl. Psychiatry 8, 44 (2018).

45. Lu, A. T. et al. Genetic architecture of epigenetic and neuronal ageing rates in human brain regions. Nat. Commun. 8, 15353 (2017).

46. Davies, M. N. et al. Functional annotation of the human brain methylome identifies tissue-specific epigenetic variation across brain and blood. Genome Biol. 13, R43 (2012).

47. Hannon, E., Lunnon, K., Schalkwyk, L. & Mill, J. Interindividual methylomic variation across blood, cortex, and cerebellum: implications for epigenetic studies of neurological and neuropsychiatric phenotypes. Epigenetics 10, 1024–1032 (2015).

48. Guintivano, J., Aryee, M. J. & Kaminsky, Z. A. A cell epigenotype specific model for the correction of brain cellular heterogeneity bias and its application to age, brain region and major depression. Epigenetics 8, 290–302 (2013).

49. Negi, S. K. & Guda, C. Global gene expression profiling of healthy human brain and its application in studying neurological disorders. Sci. Rep. 7, 897 (2017).

50. Azevedo, F. A. C. et al. Equal numbers of neuronal and nonneuronal cells make the human brain an isometrically scaled-up primate brain. J. Comp. Neurol. 513, 532–541 (2009).

51. Ma, S., Hsieh, Y.-P., Ma, J. & Lu, C. Low-input and multiplexed microfluidic assay reveals epigenomic variation across cerebellum and prefrontal cortex. Sci. Adv. 4, eaar8187 (2018).

52. Westra, H.-J. et al. Cell Specific eQTL Analysis without Sorting Cells. PLOS Genet. 11, e1005223 (2015).

53. van der Wijst, M. G. P. et al. Single-cell RNA sequencing identifies celltype-specific cis-eQTLs and co-expression QTLs. Nat. Genet. 50, 493–497 (2018).

54. Wang, J., Devlin, B. & Roeder, K. Using multiple measurements of tissue to estimate individual- and cell-type-specific gene expression via deconvolution. bioRxiv 379099 (2018). doi:10.1101/379099

55. Li, M. et al. Integrative functional genomic analysis of human brain development and neuropsychiatric risks. Science (80-.). 362, eaat7615 (2018).

56. Dobin, A. et al. STAR: ultrafast universal RNA-seq aligner. Bioinformatics 29, 15–21 (2013).

57. Liao, Y., Smyth, G. K. & Shi, W. featureCounts: an efficient general purpose program for assigning sequence reads to genomic features. Bioinformatics 30, 923–30 (2014).

58. Hansen, K. D., Irizarry, R. A. & Wu, Z. Removing technical variability in RNA-seq data using conditional quantile normalization. Biostatistics 13, 204–216 (2012).

59. Ritchie, M. E. et al. limma powers differential expression analyses for RNA-sequencing and microarray studies. Nucleic Acids Res. 43, e47–e47 (2015).

60. Law, C. W., Chen, Y., Shi, W. & Smyth, G. K. voom: precision weights unlock linear model analysis tools for RNA-seq read counts. Genome Biol. 15, R29 (2014).

61. Pearson, K. LIII. On lines and planes of closest fit to systems of points in space. London, Edinburgh, Dublin Philos. Mag. J. Sci. 2, 559–572 (1901).

62. Hotelling, H. Analysis of a complex of statistical variables into principal components. J. Educ. Psychol. 24, 417–441 (1933).

63. Leek JT, Johnson WE, Parker HS, Fertig EJ, Jaffe AE, Storey JD, Zhang Y, T. L. Bioconductor - sva. (2018). doi:10.18129/B9.bioc.sva

64. Braak, H. & Braak, E. Neuropathological stageing of Alzheimer-related changes. Acta Neuropathol. 82, 239–259 (1991).

65. Chandler, M. J. et al. A total score for the CERAD neuropsychological battery. Neurology 65, 102–106 (2005).

66. Chang, C. C. et al. Second-generation PLINK: rising to the challenge of larger and richer datasets. Gigascience 4, 7 (2015).

67. Das, S. et al. Next-generation genotype imputation service and methods. Nat. Genet. 48, 1284–1287 (2016).

68. Loh, P.-R. et al. Reference-based phasing using the Haplotype Reference Consortium panel. Nat. Genet. 48, 1443 (2016).

69. Klei, L., Kent, B. P., Melhem, N., Devlin, B. & Roeder, K. GemTools: A fast and efficient approach to estimating genetic ancestry. (2011).

70. Shabalin, A. A. Matrix eQTL: ultra fast eQTL analysis via large matrix operations. Bioinformatics 28, 1353–1358 (2012).

71. Begum, F., Ghosh, D., Tseng, G. C. & Feingold, E. Comprehensive literature review and statistical considerations for GWAS meta-analysis. Nucleic Acids Res. 40, 3777–3784 (2012).

72. O’Brien, H. & Bray, N. J. Summary statistics for expression quantitative trait loci in the developing human brain and their enrichment in neuropsychiatric disorders. (2018). doi:10.6084/m9.figshare.6881825.v1

73. Bass, A. J., Dabney, A. & Robinson Maintainer John Storey, D. D. Package ‘qvalue’ Title Q-value estimation for false discovery rate control. (2016).

