## Supplemental Tables for "Large eQTL meta-analysis reveals differing patterns between cerebral cortical and cerebellar brain regions"

### Supplementary Information for Sieberts et al.

Supplementary Table 1: Demographic and diagnosis data for samples used in the eQTL meta-analysis.

| Resource | Cohort | | Diagnosis | N | Age (years) | % Male |
|---|---|---|---|---|---|---|
| AMP-AD Consortium | ROSMAP | | AD | 138 | 88.4±2.9 | 28.3 |
| | | | Control | 82 | 83.4±5.8 | 45.1 |
| | | | Other | 353 | 86.8±4.2 | 38.0 |
| | Mayo | | AD | 80 | 82.6±7.7 | 38.8 |
| | | | Control | 71 | 82.7±8.5 | 50.7 |
| | | | Other | 111 | 77.1±7.7 | 55.0 |
| CommonMind Consortium | MSSM-Penn-Pitt | Mount Sinai (MSSM) | Schizophrenia | 110 | 74.1±12.0 | 68.2 |
| | | | Bipolar/Other | 18 | 60.9±16.1 | 38.9 |
| | | | Control | 105 | 76.5±17.8 | 49.5 |
| | | University of Pennsylvania | Schizophrenia | 47 | 81.0±6.9 | 40.4 |
| | | | Control | 25 | 67.7±17.1 | 52.0 |
| | | University of Pittsburgh | Schizophrenia | 39 | 50.0±12.4 | 82.1 |
| | | | Bipolar | 34 | 46.0±12.1 | 58.8 |
| | | | Control | 71 | 49.2±13.8 | 70.4 |
| | HBCC | | Schizophrenia | 35 | 48.6±15.1 | 71.4 |
| | | | Bipolar/Other | 47 | 43.7±15.2 | 70.2 |
| | | | Control | 67 | 43.8±16.6 | 85.1 |

Supplementary Table 2: RNA normalization and modeling used to adjust RNA-seq data prior to eQTL analysis, as well as covariates used in the eQTL model.

<table>
<tr><th colspan="2" rowspan="3">Source/Cohort</th><th rowspan="3">Brain Region (Samples)</th><th colspan="5">RNA Normalization</th><th colspan="2">eQTL Model</th></tr>
<tr><th colspan="3">Fixed effects</th><th rowspan="2">Random effect</th><th rowspan="2">Genes after Filtering</th><th rowspan="2">Number of Ancestry PCs</th><th rowspan="2">Dx</th></tr>
<tr><th>Clinical covariates</th><th>Technical covariates</th><th>Surrogate Variables</th></tr>
<tr><td rowspan="3">AMP-AD</td><td>ROSMAP</td><td>DLPFC (573)</td><td>Sex, PMI, Age of Death</td><td>Batch, RINcontinuous, PCT_CODING_BASES, PCT_INTERGENIC_BASES</td><td>21</td><td>-</td><td>15,582</td><td>3</td><td>AD, Control, Other</td></tr>
<tr><td rowspan="2">MAYO</td><td>CER (261)</td><td rowspan="2">Sex, PMI, Age of Death</td><td rowspan="2">Source, FLOWCELL, PCT_INTRONIC_BASES, RIN, PCT_INTERGENIC_BASES, PCT_CODING_BASES, PCT_RIBOSOMAL_BASES</td><td rowspan="2">19</td><td rowspan="2">Donor id</td><td rowspan="2">17,003</td><td rowspan="2">2</td><td rowspan="2">AD, Control, Other</td></tr>
<tr><td>TCX (262)</td></tr>
<tr><td rowspan="2">CommonMind</td><td>MSSM-Penn-Pitt</td><td>DLPFC (449)</td><td>Institution, Gender</td><td>RIN, IntronicRate</td><td>32</td><td>Individual</td><td>18,841</td><td>5</td><td>SCZ, Control, BP/Other</td></tr>
<tr><td>HBCC</td><td>DLPFC (149)</td><td>Gender</td><td>RIN, IntronicRate</td><td>13</td><td>-</td><td>17,250</td><td>5</td><td>SCZ, Control, BP/Other</td></tr>
</table>

Supplementary Table 3: Pathway analysis of cerebellum-specific eQTL

| GO ID | Term | Annotated Genes | Significant | Expected | *p*-value | FDR |
|---|---|---|---|---|---|---|
| GO:0007409 | axonogenesis | 138 | 42 | 23.11 | 4.00E-05 | 0.069 |
| GO:0007411 | axon guidance | 84 | 29 | 14.07 | 5.20E-05 | 0.069 |
| GO:0097485 | neuron projection guidance | 85 | 29 | 14.24 | 6.60E-05 | 0.069 |
| GO:0051962 | positive regulation of nervous system development | 173 | 49 | 28.97 | 7.40E-05 | 0.069 |
| GO:0061564 | axon development | 153 | 44 | 25.62 | 1.2E-04 | 0.074 |
| GO:0048699 | generation of neurons | 447 | 104 | 74.86 | 1.3E-04 | 0.074 |
| GO:0048667 | cell morphogenesis involved in neuron differentiation | 177 | 49 | 29.64 | 1.4E-04 | 0.074 |
| GO:0048812 | neuron projection morphogenesis | 192 | 52 | 32.16 | 1.6E-04 | 0.074 |
| GO:0048468 | cell development | 627 | 137 | 105.01 | 2.4E-04 | 0.096 |
| GO:0048522 | positive regulation of cellular process | 1499 | 294 | 251.05 | 2.8E-04 | 0.096 |
| GO:0010605 | negative regulation of macromolecule metabolic process | 729 | 155 | 122.09 | 3.5E-04 | 0.096 |
| GO:0051149 | positive regulation of muscle cell differentiation | 29 | 13 | 4.86 | 3.6E-04 | 0.096 |
| GO:0030182 | neuron differentiation | 403 | 93 | 67.49 | 4.0E-04 | 0.096 |
| GO:0022008 | neurogenesis | 476 | 107 | 79.72 | 4.2E-04 | 0.096 |
| GO:0120039 | plasma membrane bounded cell projection morphogenesis | 199 | 52 | 33.33 | 4.2E-04 | 0.096 |
| GO:0021953 | central nervous system neuron differentiation | 48 | 18 | 8.04 | 4.3E-04 | 0.096 |
| GO:0051172 | negative regulation of nitrogen compound metabolic processes | 615 | 133 | 103 | 4.7E-04 | 0.096 |
| GO:0048858 | cell projection morphogenesis | 200 | 52 | 33.5 | 4.8E-04 | 0.096 |
| GO:0032990 | cell part morphogenesis | 205 | 53 | 34.33 | 4.9E-04 | 0.096 |
